# Identification of ESBL-producing enterobacterales from vegetable plants in a rural area of Madagascar: insights into AMR transmission at the human-animal-environment-plant interface

**DOI:** 10.1101/2025.04.06.647499

**Authors:** Adrien Rieux, Mamitina Alain Noah Rabenandrasana

**Author notes:** <u>Author emails:</u>.

## Abstract

Extended-spectrum b-lactamases (ESBL)-producing enterobacterales are considered a key indicator for antimicrobial resistance (AMR) epidemiological surveillance in animal, human and environment compartments. In this study we aim to investigate the presence and genetic diversity of ESBL-producing enterobacterales on vegetable plants. We isolated beta-lactam resistant enterobacterales from several vegetable plants and sequenced their whole genome. Utilizing cutting-edge genomic and phylogenetic methods, we sought to *i)* characterize their resistome and plasmidome, *ii)* investigate the genetic structure of the plant-isolated strains and *iii)* determine their relationships with strains from other reservoirs. Among the 22 strains collected from vegetable plants, 6 showed resistance to beta-lactam antibiotics, with 5 of them identified as ESBL producers. Resistome analysis indicated the presence of multidrug-resistant (MDR) strains containing multiple antibiotic resistance genes (ARGs). Importantly, no host-specific lineages were identified among the plant-isolated ESBL-producing *E. coli* (ESBL-Ec). Instead, these strains exhibited genetic and epidemiological connections with strains isolated from animals, humans, and the environment, suggesting transfer of ESBL-Ec between plants and other sources in rural Madagascar. These findings suggest that vegetable plants are contaminated as a result of human activities, posing a potential risk of human and animal exposure to antibiotic-resistant bacteria and genes.

## Introduction

Extended-spectrum *β*-lactamase (ESBL)-producing Gram-negative bacteria are a leading cause of human and animal infection, being categorized as critical priority pathogens by the World Health Organization^1^. In this context, antimicrobial resistance (AMR) is recognized as a major One Health challenge because of the rapid emergence and dissemination of resistant bacteria and genes among humans, animals and the environment^2^. Aside from assessing AMR prevalence, there is an increasing need to better understand behaviors, customs and practices that drive the evolution and transmission of resistance, both globally and in low-resource settings in which the AMR threat is of particular concern^3^. In this context, the use of antibiotics in agriculture has historically been described as a major contributor of AMR in humans due to a large variety of potential dissemination pathways, with most of them involving the influence of livestock^4^. The role of livestock in the emergence of AMR was shown to be multifaceted, involving both direct transmission to humans and environmental contamination^5^. In contrast, little is known about the existence and prevalence of antimicrobial resistant bacteria and resistance genes (ARB & ARG, respectively) within the cultivated plant reservoir, despite the fact that such food is often consumed raw, hence increasing the risk of transmission to humans^6^. Herein, we aimed to characterize the prevalence and genetic diversity of beta-lactam resistant enterobacterales isolated from vegetable plants in a rural area of Madagascar. To assess the genetic and epidemiological relationships between strains isolated from plants and other reservoirs, we integrated data from a cross-sectional population-based recent study performed within the same area on animals, humans and the environment^7^.

### Experimental procedures

#### Sampling, bacterial culture and sequencing

In September 2018, 22 vegetable plants (spinach, lettuce, tomato & cabbage) were sampled within 6 backyard gardens in Andoharanofotsy, Madagascar (FigS1). Sampling was performed using sterile equipment. Plant leaves were transported back to the laboratory in individual envelopes before being finely cut and soaked in LB broth for 24 h at 35 ± 2°C with continuous shaking. 100 μL of the enriched suspension was directly streaked onto selective chromogenic agar plates (CHROMagar ESBL; CHROMagar, Paris, France) and incubated overnight at 35 ± 2°C under aerobic conditions. All presumptive ESBL-producer morphotypes were subcultured individually on LB agar plates.

*In vitro* antimicrobial susceptibility testing was performed on one isolate according to the standard disc methods described in the 2024 ‘Comité de l’Antibiogramme de la Société Française de Microbiologie’ (CASFM)-EUCAST guidelines^8^. The presence of ESBL enzymes was confirmed by synergy of cefotaxime, ceftazidime and cefepime with amoxicillin/clavulanate or ticarcillin/clavulanate.

One bacterial isolate per sample was randomly selected for genetic analysis. DNA extraction was performed using the Cador Pathogen Extraction Kit (INDICAL Bioscience) from 5 mL of liquid cultures grown overnight at 37°C in LB broth medium. Library preparation was performed using the Nextera XT DNA Library Preparation Kit (Illumina, San Diego, CA, USA) and sequencing was performed on a NextSeq 500 platform (Illumina) using 2 × 150 bp runs.

#### Core genome analyses

Raw reads were trimmed with Trimmomatic^9^ to remove adapters and low-quality sequences before being *de novo* assembled using the Unicycler pipeline^10^ (default parameters). Taxonomic assignment was performed on assembled bacterial genomes both by computing ANI using FastANI^11^ and by running Kraken on the microbial reference database^12^. In order to investigate the phylogenetic structure of the plant-isolated ESBL genomes within a global One-health framework, we aligned the bacterial genomes generated within the course of this study with a public dataset of 510 ESBL strains also isolated in 2018 from human, animal and water in the same rural area of Madagascar^7^. Genomes were aligned using parsnp^13^ and regions acquired via horizontal gene transfers were detected using Gubbins^14^. From the recombination-free SNP alignment, a maximum likelihood phylogeny was constructed using RAxML^15^ with a rapid bootstrap analysis, general time-reversible model of evolution with a four rate categories γ distribution (GTRGAMMA) and 1000 iterations. Transmission clusters were inferred from the reconstructed phylogeny using the dedicated Phydelity tool^16^.

#### Genotyping, mobilome and resistome analysis

Sequence types and phylogroups were estimated *in silico* using staram^17^ and ClermonTyping^18^ softwares, respectively. Staramr was also used to scan bacterial genome contigs against the ResFinder, PointFinder, and PlasmidFinder databases to detect the presence of plasmids and antimicrobial resistance genes. RFPlasmid^19^ was used to predict whether the detected ARG were located on plasmids or chromosomes.

## Results

### Prevalence of beta-lactam resistant and ESBL producers enterobacterales in vegetable plants

We identified beta-lactam resistant enterobacterales from 6 out of 22 collected plants (average prevalence of 27%, Fig.1A). Positive samples were isolated from 5 out of the 6 sampled backyard gardens and originated from lettuce (n=2), spinach (n=2), cabbage (n=1) and tomato (n=1) (Fig. 1B).

**Fig 1:**
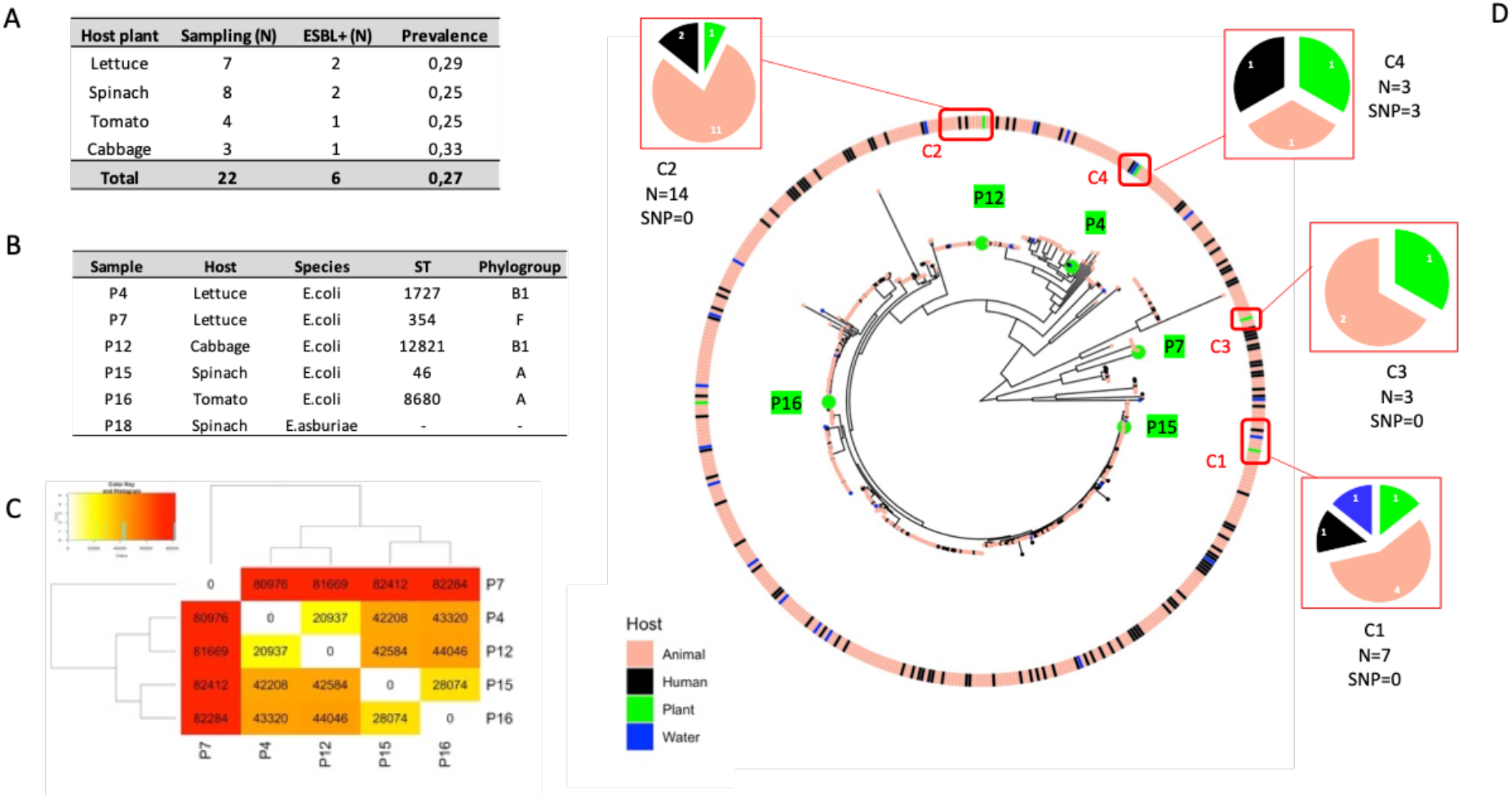
A) Sampled plants and prevalence of isolated beta-lactam resistant enterobacterales. B) Taxonomic assignment of isolated enterobacterales. C) Pairwise SNP number between the five ESBL-Ec. D) Phylogenetic maximum likelihood tree of 515 ESBL-Ec genomes built from 329817 core and non-recombinant SNPs. Hosts for each isolate (human, animal, plant, water) are depicted using different colors, with plant isolates highlighted in green. Transmission clusters and their composition are indicated with red frames and pie charts, respectively. The number of isolates in each cluster (N) and their average SNP count (SNP) are shown below.

**Fig 2:**
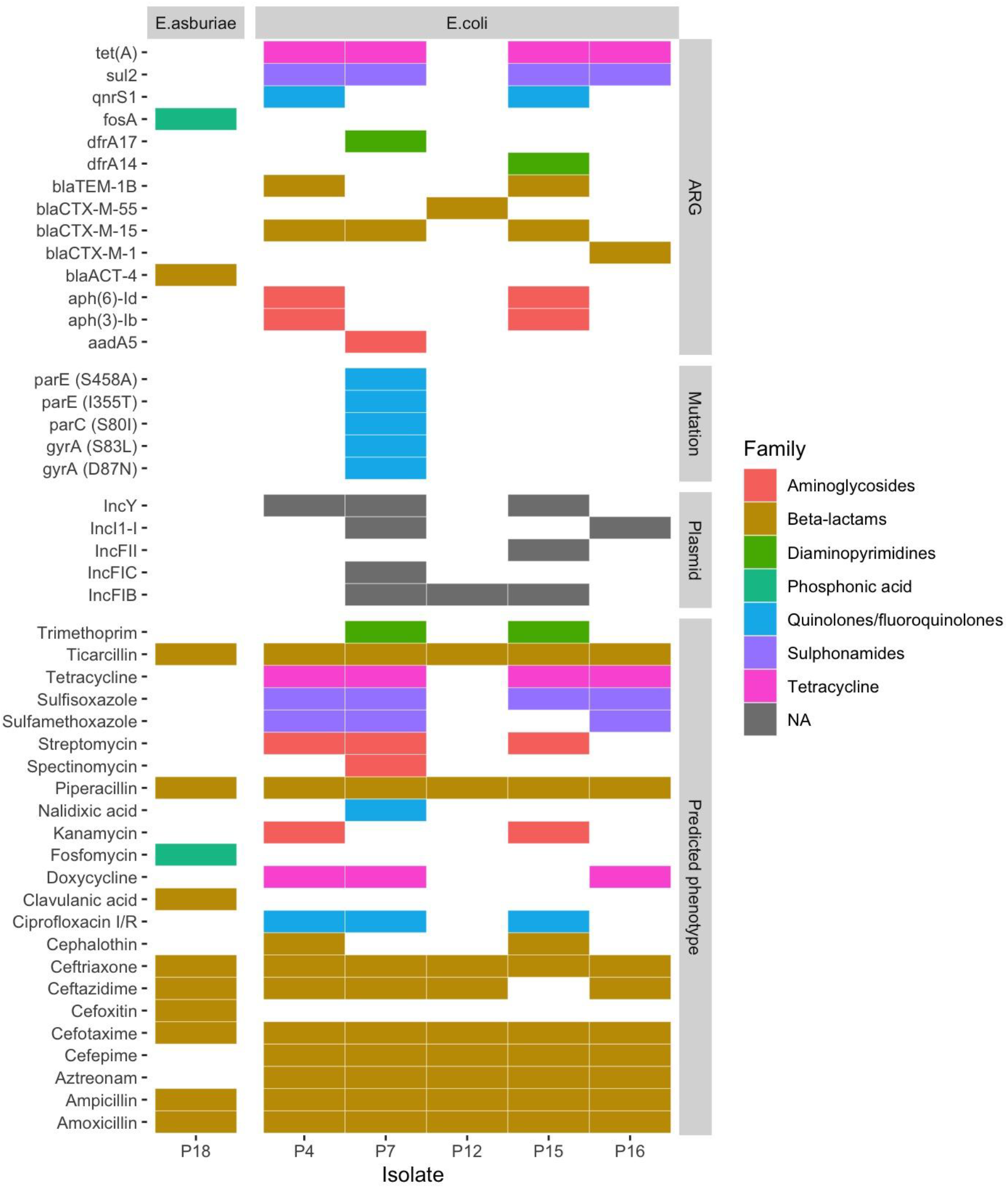
Distribution of AMR genotypes and predicted resistance phenotypes among the six plant-isolated beta-lactam resistant enterobacterales strains. Filled boxes indicate the presence of ARGs, point mutations, plasmids, or a predicted antibiotic-resistant phenotype, with colors corresponding to different antibiotic families.

*In vitro* antimicrobial susceptibility testing yielded variable results according to the strains and antibiotics tested (Fig S2). All 6 strains were sensitive to Ertapenem & Imipenem, resistant to Amoxicillin, Ceftazidim & Cefotaxim and showed variable sensitivity patterns towards the other tested antibiotics. Five strains (P4, P7, P12, P15 & P16) passed the synergy test and were hence considered as ESBL producers. Strain P18, although resistant to cefotaxime, ceftazidime, ticarcillin/clavulanate & amoxicillin/clavulanate did not produce any ESBL.

### Bacterial sequencing

Sequencing generated around 8 million paired-end reads with a majority (99.90%–99.96%) of bases scoring Q30 and above. *Denovo* assembly statistics are given in Table S1.

### Taxonomic assignment, mobilome and resistome

Out of the 6 beta-lactam resistant enterobacterales, 5 were assigned to *Escherichia coli* and 1 to *Enterobacter asburiae* species. The five E.coli, all ESBL-producers, were assigned to three different phylogroups (A; n=2, B1; n=2 and F; n=1) and 5 STs (1727, 354, 12821, 46 & 880), respectively.

A total of 14 different ARGs, 5 point mutations and 5 plasmids were identified among the six plant beta-lactam resistant enterobacterales (Figure 1 & Table S2). In total, the six isolated strains harbored ARGs against seven different antibiotic families. The five ESBL-Ec strains were predicted to be resistant to at least three antibiotic families and hence considered as multiple drug resistant (MDR). All six strains were predicted to be resistant to several antibiotics from the Beta-lactams family. When comparing resistance patterns predicted *in silico* with those measured *in vitro*, we observed consistent results for 94% (74/78) of the strains/antibiotics tested (Figure S3).

*Bla*_CTX-M-15_ was the most frequently detected ESBL-gene and a total of 5 different plasmids (IncY, IncFIB, IncFIC, IncFII & IncI1-I) were shown to segregate within the sampled strains. Detected ARGs were predicted to be carried either by chromosomes or plasmids (Table S2) except for *Bla*_CTX-M-15_ which was detected both on a chromosome (strain P7) and a plasmid (strain P4). Most detected ARGs displayed very high nucleotide identity rates with reference genes (Table S2).

### Genomic diversity and phylogenetic structure

As illustrated by their assignment to unique STs, the five ESBL-Ec strains isolated from plants displayed high levels of genomic diversity with a mean pairwise SNP number = 54851 (Fig1C). In order to decipher their genetic relationships with strains isolated from other reservoirs, we built a phylogenetic tree using a dataset composed of previously published ESBL-Ec genomic resources^7^ (Total of 329817 SNPs in between 515 ESBL-Ec strains). Interestingly, plant ESBL-Ec strains did not cluster together within the phylogenetic tree but were instead intermixed within other human/animal/environment-isolated strains (Fig1D). Furthermore, among the five plant ESBL-Ec strains, four were assigned to transmission clusters with other strains originating from other compartments (Fig1D&TableS3). Strain P16, isolated from tomato leaf, was not assigned to any transmission cluster.

## Discussion

In this study, we report for the first time the presence of ESBL-producing enterobacterales from vegetable plants sampled in Madagascar. In this regard, the occurrence of ESBL producers in various crops and fresh vegetables have previously been described in other countries^6,20,21^. Interestingly, members of the enterobacterales order have previously been shown to exhibit both an endophytic (i.e. colonize internal tissues) and epiphytic (i.e. live and multiply on the outside of aerial surfaces) lifestyle on vegetable plants^20^. Unfortunately, the methodology used in this study did not allow for a distinction between the two lifestyle modes. Amongst the five E-coli strains recovered, isolate P15 sampled from spinach was assigned to ST46 which encompass pathogenic strains responsible for septicaemia, diarrhoea, meningitis, and urinary tract infections^22^. In addition, P18, also isolated from spinach, was assigned to *E. asburiae*, a bacterial species previously shown to be associated with humans and animals while also causing disease on plants such as rice and radish^23,24^.

Determinants of antimicrobial resistance and plasmids were *in silico* detected from the genomic data. Amongst the detected genes providing resistance to Beta-lactams, *Bla*_CTX-M-15_ and *Bla*_CTX-M-55_ are both members of the CTX-M-1 group, considered as one of the most predominant enzymes in the widespread dissemination of ESBLs^25^. Interestingly, most of those ESBL-genes were *i)* estimated to be plasmid-mediated and *ii)* previously detected within different compartments in Madagascar^7,26^, highlighting their relevance to One Health concerns. In this context, a recent study revealed frequent and multiple transmission events between ESBL-Ec strains isolated from humans, animals and the environment (drinking water) in a suburban rural area of Antananarivo, Madagascar. Through a dedicated phylogenetic analysis, we demonstrated that the newly isolated ESBL-Ec strains from vegetable plants were interspersed within the genomic diversity of strains from the other compartments, as illustrated by the inference of shared transmission clusters. The lack of genomic diversity within the detected clusters supports a likely recent epidemiological connection between strains from vegetable plants and other reservoirs.

Although, the origins of ESBL-producing enterobacterales found in this study on vegetable plants were not investigated, they could originate from human (as sewage), animal (manure), and/or environmental (such as contaminated soil and irrigation water) sources that come into contact with crops^27,28^. Therefore, our pilot results suggest that the consumption of raw vegetables should be seen as a public health problem and a potential risk of human exposure to antibiotic-resistant bacteria and/or their resistance genes. Future studies performed on a higher number of samples are required to decipher more accurately the drivers and pathways of ARB & ARG transmission between plants and other compartments, in order to gather data for risk assessment.

## Author contributions

A.R designed the study and performed field-sampling. M.A.N.R. performed lab work and assisted the sequencing work. A.R. & M.A.N.R both performed genomic analyses and wrote the manuscript.

## Acknowledgements

We thank Noellie Gay/the Andoharanofotsy community (mayors, heads of fokontanys) for help with sampling and the staff of the Plateforme de Microbiologie Mutualisée (P2M) at Institut Pasteur Paris for their help with whole-genome sequencing. We are also grateful to Tania Crucitti and Olivier Pruvost for valuable comments and discussions. Computational work was performed on the MESO@LR-Platform at the University of Montpellier (https://hal.umontpellier.fr/MESO).

## Data availability

Both raw reads and assembled genomes data are available from NCBI under the accession number listed in TableS1.

## Supplementary data

- Fig S1: Sampling sites in Andoharanofotsy, Madagascar
- Fig S2: In vitro antimicrobial susceptibility results.
- Fig S3: Comparison between measured and predicted resistant phenotypes
- Table S1: Genomic data generated for the 6 plant-isolated enterobacterales
- Table S2: Resistome and plasmidome of the plant-isolated enterobacterales
- Table S3: Composition of ESBL-Ec transmission clusters (estimated with Phidelity) involving plant-isolated strains.

