## Supplementals for "Identification of ESBL-producing enterobacterales from vegetable plants in a rural area of Madagascar: insights into AMR transmission at the human-animal-environment-plant interface"

| Sample | Site | GPS coord | Vegetable plant | ESBL |
|---|---|---|---|---|
| P1 | S1 | S 18,99408 E 047,54015 | spinach | Negative |
| P2 | S1 | S 18,99408 E 047,54015 | lettuce | Negative |
| P3 | S1 | S 18,99408 E 047,54015 | lettuce | Negative |
| P4 | S1 | S 18,99408 E 047,54015 | lettuce | Positive |
| P5 | S2 | S 18,99916 E 047,53862 | spinach | Negative |
| P6 | S2 | S 18,99916 E 047,53862 | tomato | Negative |
| P7 | S2 | S 18,99916 E 047,53862 | lettuce | Positive |
| P8 | S3 | S 18,99064 E 047,53586 | spinach | Negative |
| P9 | S3 | S 18,99064 E 047,53586 | cabbage | Negative |
| P10 | S3 | S 18,99064 E 047,53586 | lettuce | Negative |
| P11 | S3 | S 18,99064 E 047,53586 | lettuce | Negative |
| P12 | S3 | S 18,99064 E 047,53586 | cabbage | Positive |
| P13 | S4 | S 18,97230 E 047,42816 | cabbage | Negative |
| P14 | S4 | S 18,97230 E 047,42816 | spinach | Negative |
| P15 | S4 | S 18,97230 E 047,42816 | spinach | Positive |
| P16 | S4 | S 18,97230 E 047,42816 | tomato | Positive |
| P17 | S5 | S 18,97200 E 047,52409 | lettuce | Negative |
| P18 | S5 | S 18,97200 E 047,52409 | spinach | Positive |
| P19 | S6 | S 18,99850 E 047,54042 | spinach | Negative |
| P20 | S6 | S 18,99850 E 047,54042 | spinach | Negative |
| P21 | S6 | S 18,99850 E 047,54042 | tomato | Negative |
| P22 | S6 | S 18,99850 E 047,54042 | tomato | Negative |

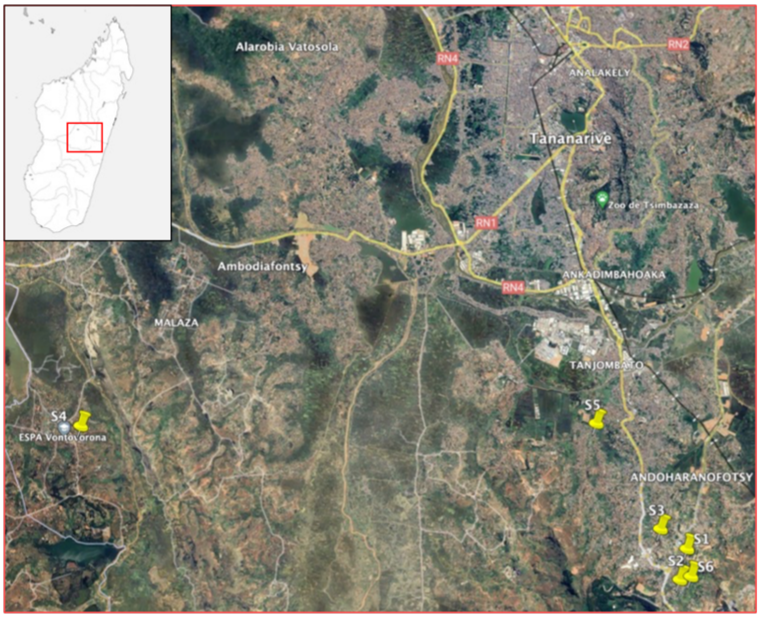


### Fig S1: Sampling sites in Andoharanofotsy, Madagascar

| ID | P4 | P7 | P12 | P15 | P16 | P18 |
|---|---|---|---|---|---|---|
| Amoxicillin | 6 | 6 | 6 | 6 | 6 | 6 |
| Amoxicillin-clavulanate | 20 | 22 | 21 | 20 | 22 | 6 |
| Aztreonam | 18 | 19 | 16 | 19 | 18 | 30 |
| Cefepim | 16 | 18 | 18 | 18 | 19 | 40 |
| Cefotaxim | 6 | 6 | 6 | 9 | 6 | 6 |
| Cefoxitin | 21 | 21 | 26 | 27 | 22 | 6 |
| Ceftazidim | 18 | 16 | 15 | 18 | 17 | 14 |
| Ciprofloxacin | 29 | 10 | 31 | 31 | 32 | 46 |
| Ertapenem | 25 | 26 | 26 | 25 | 29 | 28 |
| Imipenem | 30 | 30 | 32 | 29 | 31 | 30 |
| Nalidixic Acid | 25 | 11 | 23 | 23 | 24 | 38 |
| Ticarcillin-clavulanate | 23 | 24 | 25 | 24 | 24 | 6 |
| Trimethoprim-Sulfamethoxazole | 21 | 10 | 23 | 11 | 22 | 30 |
| Synergy_test_ESBL | Positive | Positive | Positive | Positive | Positive | Negative |

| | |
|---|---|
| (red) | *In vitro* measured resistant phenotype |
| (green) | *In vitro* measured sensitive phenotype |

Values indicate diameters of bacterial growth inhibition measured on agar plates (in mm)

**Fig S2: *In vitro* antimicrobial susceptibility results.**

| ID | P4 | P7 | P12 | P15 | P16 | P18 |
|---|---|---|---|---|---|---|
| Amoxicillin | R | R | R | R | R | R |
| Amoxicillin-clavulanate | S | S | S | S | S | R |
| Aztreonam | R | R | R | R | R | S |
| Cefepim | R | R | R | R | R | S |
| Cefotaxim | R | R | R | R | R | R |
| Cefoxitin | S | S | S | S | S | R |
| Ceftazidim | R | R | R | S | R | R |
| Ciprofloxacin | R | R | S | R | S | S |
| Ertapenem | S | S | S | S | S | S |
| Imipenem | S | S | S | S | S | S |
| Nalidixic Acid | S | R | S | S | S | S |
| Ticarcillin-clavulanate | S | S | S | S | S | R |
| Trimethoprim-Sulfamethoxazole | S | R | S | S | S | S |

| | |
|---|---|
| | *In vitro* -measured resistant phenotype |
| | *In vitro* -measured sensitive phenotype |
| R | In siico-predicted resistant phenotype |
| S | In siico-predicted sensitive phenotype |

Discrepancy between measured & predicted patterns

**Fig S3: Comparison between measured and predicted resistant phenotypes**

| Sample | P4 | P7 | P12 | P15 | P16 | P18 |
|---|---|---|---|---|---|---|
| Vegetable plant | Salad | Salad | Cabbage | Spinach | Tomato | Spinach |
| Scientific name | *Lactuca sativa* | *Lactuca sativa* | *Brassica oleracea* | *Spinacia oleracea* | *Solanum lycopersicum* | *Spinacia oleracea* |
| Taxonomic assignment | *Esherichia coli* | *Esherichia coli* | *Esherichia coli* | *Esherichia coli* | *Esherichia coli* | *Enterobacter cloacae* |
| ST | 1727 | 354 | 12821 | 46 | 8680 | - |
| Phylogroup | B1 | F | B1 | A | A | - |
| Genome size (Gb) | 4,64 | 5,09 | 4,76 | 4,63 | 4,48 | 4,68 |
| Number of contigs | 107 | 250 | 171 | 174 | 191 | 42 |
| N50 value | 72520 | 49481 | 51290 | 55944 | 42774 | 171452 |
| Bioproject | PRJNA1147432 | | | | | |
| BioSample | SAMN43154107 | SAMN43154108 | SAMN43154109 | SAMN43154110 | SAMN43154111 | SAMN43154112 |
| SRA (raw reads) | SRR30230049 | SRR30230048 | SRR30230047 | SRR30230046 | SRR30230045 | SRR30230044 |
| Assembled genome | JBGMDQ000000000 | JBGMDP000000000 | JBGMDO000000000 | JBGMDN000000000 | JBGMDM000000000 | JBGMDL000000000 |

**Table S1: Genomic data generated for the 6 plant-isolated enterobacterales**

| Isolate ID | Data | Data Type | Predicted Phenotype | Genetic support (probability) | %Identity | %Overlap | HSP Length/Total Length | Contig | Start | End | Accession |
|---|---|---|---|---|---|---|---|---|---|---|---|
| **P12** | IncFIB | Plasmid | | Plasmid (0.71) | 100 | 100 | 560/560 | 149 | 464 | 1023 | CU638872 |
| **P12** | blaCTX-M-55 | Resistance | Amoxicillin, Ampicillin, Aztreonam, Cefepime, Cefotaxime, Ceftriaxone, Ceftazidime, Piperacillin, Ticarcillin | Chromosome (1) | 100 | 100 | 876/876 | 152 | 1144 | 2019 | DQ810789 |
| **P15** | IncFIB(AP001918) | Plasmid | | Plasmid (0.87) | 99,12 | 100 | 682/682 | 114 | 1835 | 2516 | AP001918 |
| **P15** | IncFII | Plasmid | | Plasmid (0.99) | 96,56 | 100 | 261/261 | 42 | 1662 | 1923 | AY458016 |
| **P15** | IncY | Plasmid | | Plasmid (0.90) | 98,82 | 100 | 765/765 | 50 | 27922 | 28686 | K02380 |
| **P15** | aph(3'')-Ib | Resistance | Streptomycin | Plasmid (0.95) | 100 | 100 | 804/804 | 36 | 35754 | 34951 | AF321551 |
| **P15** | aph(6)-Id | Resistance | Streptomycin, kanamycin | Plasmid (0.95) | 100 | 100 | 837/837 | 36 | 34951 | 34115 | M28829 |
| **P15** | blaCTX-M-15 | Resistance | Amoxicillin, Ampicillin, Aztreonam, Cefepime, Cefotaxime, Ceftazidime, Ceftriaxone, Piperacillin, Ticarcillin | Plasmid (0.95) | 100 | 100 | 876/876 | 36 | 29712 | 28837 | AY044436 |
| **P15** | blaTEM-1B | Resistance | Amoxicillin, Ampicillin, Cephalothin, Piperacillin, Ticarcillin | Plasmid (0.95) | 100 | 100 | 861/861 | 36 | 32534 | 33394 | AY458016 |
| **P15** | dfrA14 | Resistance | Trimethoprim | Plasmid (0.72) | 100 | 100 | 474/474 | 142 | 1162 | 1635 | KF921535 |
| **P15** | qnrS1 | Resistance | Ciprofloxacin | Plasmid (0.95) | 100 | 100 | 657/657 | 36 | 24196 | 23540 | AB187515 |
| **P15** | sul2 | Resistance | Sulfamethoxazole, Sulfisoxazole | Plasmid (0.95) | 100 | 100 | 816/816 | 36 | 36630 | 35815 | AY034138 |
| **P15** | tet(A) | Resistance | Doxycycline, Tetracycline | Plasmid (0.91) | 100 | 100 | 1200/1200 | 124 | 1054 | 2253 | AJ517790 |
| **P16** | IncI1-I(Alpha) | Plasmid | | Plasmid (0.93) | 99,3 | 100 | 142/142 | 63 | 5034 | 5175 | AP005147 |
| **P16** | blaCTX-M-1 | Resistance | Amoxicillin, Ampicillin, Aztreonam, Cefepime, Cefotaxime, Ceftazidime, Ceftriaxone, Piperacillin, Ticarcillin | Plasmid (0.99) | 100 | 100 | 876/876 | 8 | 13545 | 14420 | DQ915955 |
| **P16** | sul2 | Resistance | Sulfamethoxazole, Sulfisoxazole | Plasmid (0.93) | 100 | 100 | 816/816 | 63 | 16389 | 17204 | AY034138 |
| **P16** | tet(A) | Resistance | Doxycycline, Tetracycline | Plasmid (0.93) | 100 | 100 | 1200/1200 | 63 | 12731 | 11532 | AJ517790 |
| **P18** | blaACT-4 | Resistance | Amoxicillin, Amoxicillin+Clavulanic acid, Ampicillin, Ampicillin+Clavulanic acid, Cefotaxime, Ceftriaxone Cefoxitin, Ceftazidime, Piperacillin, Piperacillin+Tazobactam, Ticarcillin, Ticarcillin+Clavulanic acid | Chromosome (0.98) | 96,34 | 100 | 1146/1146 | 14 | 17359 | 16214 | AJ311172 |
| **P18** | fosA | Resistance | Fosfomycin | Plasmid (0.99) | 95,3 | 100 | 426/426 | 2 | 311450 | 311025 | AEXB01000013 |
| **P4** | IncY | Plasmid | | Plasmid (0.82) | 98,82 | 100 | 765/765 | 60 | 23140 | 23904 | K02380 |
| **P4** | aph(3'')-Ib | Resistance | Streptomycin | Plasmid (0.95) | 100 | 100 | 804/804 | 81 | 2278 | 1475 | AF321551 |
| **P4** | aph(6)-Id | Resistance | Streptomycin, kanamycin | Plasmid (0.95) | 100 | 100 | 837/837 | 81 | 1475 | 639 | M28829 |
| **P4** | blaCTX-M-15 | Resistance | Amoxicillin, Ampicillin, Aztreonam, Cefepime, Cefotaxime, Ceftazidime, Ceftriaxone, Piperacillin, Ticarcillin | Plasmid (0.89) | 100 | 100 | 876/876 | 78 | 6292 | 5417 | AY044436 |
| **P4** | blaTEM-1B | Resistance | Amoxicillin, Ampicillin, Cephalothin, Piperacillin, Ticarcillin | Plasmid (0.89) | 100 | 100 | 861/861 | 78 | 9114 | 9946 | AY458016 |
| **P4** | qnrS1 | Resistance | Ciprofloxacin | Plasmid (0.89) | 100 | 100 | 657/657 | 78 | 776 | 120 | AB187515 |
| **P4** | sul2 | Resistance | Sulfamethoxazole, Sulfisoxazole | Plasmid (0.95) | 100 | 100 | 816/816 | 81 | 3154 | 2339 | AY034138 |
| **P4** | tet(A) | Resistance | Doxycycline, Tetracycline | Plasmid (0.72) | 100 | 100 | 1200/1200 | 87 | 2848 | 1649 | AJ517790 |
| **P7** | IncFIB(AP001918) | Plasmid | | Plasmid (0.73) | 98,53 | 100 | 682/682 | 165 | 2629 | 1948 | AP001918 |
| **P7** | IncFIC(FII) | Plasmid | | Plasmid (0.93) | 95,79 | 100 | 499/499 | 83 | 13564 | 13068 | AP001918 |
| **P7** | IncI1-I(Alpha) | Plasmid | | Plasmid (0.75) | 94,93 | 97,18 | 138/142 | 182 | 2237 | 2102 | AP005147 |
| **P7** | IncY | Plasmid | | Plasmid (0.72) | 99,08 | 100 | 765/765 | 106 | 3159 | 3923 | K02380 |
| **P7** | aadA5 | Resistance | Spectinomycin, Streptomycin | Plasmid (0.97) | 100 | 100 | 789/789 | 80 | 926 | 138 | AF137361 |
| **P7** | blaCTX-M-15 | Resistance | Amoxicillin, Ampicillin, Aztreonam, Cefepime, Cefotaxime, Ceftazidime, Ceftriaxone, Piperacillin, Ticarcillin | Chromosome (0.91) | 100 | 100 | 876/876 | 20 | 4279 | 3404 | AY044436 |
| **P7** | dfrA17 | Resistance | Trimethoprim | Plasmid (0.97) | 100 | 100 | 474/474 | 80 | 1530 | 1057 | FJ460238 |
| **P7** | gyrA (D87N) | Resistance | Nalidixic acid;Nalidixic acid,Ciprofloxacin | Chromosome (1) | 97,83 | 100 | 2628/2628 | 53 | 26230 | 23603 | |
| **P7** | gyrA (S83L) | Resistance | Nalidixic acid,Ciprofloxacin | Chromosome (1) | 97,83 | 100 | 2628/2628 | 53 | 26230 | 23603 | |
| **P7** | parC (S80I) | Resistance | Nalidixic acid,Ciprofloxacin | Chromosome (1) | 98,05 | 100 | 2259/2259 | 24 | 21427 | 23685 | |
| **P7** | parE (I355T) | Resistance | Nalidixic acid,Ciprofloxacin | Chromosome (1) | 96,4 | 100 | 1893/1893 | 24 | 6260 | 8148 | |
| **P7** | parE (S458A) | Resistance | Nalidixic acid,Ciprofloxacin | Chromosome (1) | 96,4 | 100 | 1893/1893 | 24 | 6260 | 8148 | |
| **P7** | sul2 | Resistance | Sulfamethoxazole, Sulfisoxazole | Plasmid (0.92) | 100 | 100 | 816/816 | 148 | 1444 | 2259 | AY034138 |
| **P7** | tet(A) | Resistance | Doxycycline, Tetracycline | Plasmid (0.97) | 100 | 100 | 1200/1200 | 80 | 8048 | 9247 | AJ517790 |

**Table S2: Resistome and plasmidome of the plant-isolated enterobacterales**

| Cluster ID | Sample ID | Host | Host details | Mean pairwise SNP |
|---|---|---|---|---|
| C1 | P15 | Plant | Spinach | 0 |
| | DI12_1 | Animal | Turkey | |
| | DI8_1 | Animal | Turkey | |
| | E10_1 | Water | Water | |
| | H294_1 | Human | Human | |
| | PG47_1 | Animal | Poultry | |
| C2 | P12 | Plant | Cabbage | 0 |
| | BV23_1 | Animal | Cattle | |
| | CH10_1 | Animal | Dog | |
| | CH21_1 | Animal | Dog | |
| | H242_1 | Human | Human | |
| | H283_1 | Human | Human | |
| | OI31_1 | Animal | Goose | |
| | OI37_1 | Animal | Goose | |
| | PC30_1 | Animal | Pig | |
| | PC51_1 | Animal | Pig | |
| | PC77_1 | Animal | Pig | |
| | PG137_1 | Animal | Poultry | |
| | PG366_1 | Animal | Poultry | |
| | PP12_1 | Animal | Poultry | |
| C3 | P7 | Plant | Salad | 0 |
| | CH89_1 | Animal | Dog | |
| | OI47_1 | Animal | Goose | |
| C4 | P4 | Plant | Salad | 3 |
| | E35_1 | Water | Water | |
| | H7_1 | Human | Human | |

**Table S3: Composition of ESBL-Ec transmission clusters (estimated with Phidelity) involving plant-isolated strains.**
